# Exploratory study of factors associated with human brucellosis in mainland China based on time-series-cross-section data from 2005 to 2016

**DOI:** 10.1101/472340

**Authors:** Yun Lin, Tao Zhang, Xingyu Zhang

## Abstract

**Objective:** Many studies focused on reasons behind the increasing incidence and the spread of human brucellosis in mainland China, yet most of them lacked comprehensive consideration with quantitative evidence. Hence, this study aimed to further investigate the epidemic mechanism and associated factors of human brucellosis in China so as to provide suggestions on more effective countermeasures.

**Methods:** Data of human brucellosis incidence and some associated factors in economy, animal husbandry, transportation and health were collected at provincial level from 2005-2016. Time series plot and cluster analysis were first used to visualize incidence levels and categorize provinces based on their incidence level and epidemic trend of human brucellosis. Furthermore, according to the characteristics of data, the dynamic panel data model in combination with supervised principal component analysis was proposed to explore the effects of associated factors on human brucellosis.

**Results:** ① The incidence rate of human brucellosis has increased threefold (from 1.41 in 2005 to 4.22 in 2016) in mainland China. Incidence rates in the north have always been higher than those in the south, but the latter also experienced an upward trend especially in the recent five years. ② The 31 provinces of mainland China were categorized into three clusters, and each cluster had its own characteristics of incidence level and epidemic trend. ③ Public health expenditure and rural medical expenditure proportion were potential protective factors of human brucellosis, with attribute risks of −0.74 and −1.04 respectively. Other factors (such as amount of sheep, total length of highways, etc.) exhibited relatively trivial effects.

**Conclusions:** The epidemic status of human brucellosis has changed in both spatial and temporal dimensions in recent years. Apart from those traditional control measures, more attention should be paid to the improvement of medical healthcare especially in rural areas in order to strengthen the control effect.

## Introduction

Brucellosis is a highly contagious zoonotic disease mainly caused by unpasteurized milk or undercooked meat products from infected animals. Direct contacts with ill animals can also cause brucellosis infection. On a global scale, brucellosis was and still is an important zoonosis across the world [1–3]. In China [4], it has been listed as the class B notifiable infectious disease since 2005 as well as one of the most serious types of class B diseases among those listed in the *Detailed Rules for the Implementation of the Regulations on Livestock and Poultry Epidemic Prevention.* The incidence of human brucellosis in mainland China decreased during the 1980-1990s but then rose steadily since 1995 till the 2014 peak. Apart from the temporal trend, the epidemic of human brucellosis in mainland China also had some spatial characteristics. In the past [5–6], brucellosis (both among human and animals) was most severe in the northeast area in China possibly due to [7] that the massive number of pastures increased local residents’ risks of exposure to infected animals or their products. However, some recent studies [6] showed that human brucellosis epidemics were spreading from traditional high-incidence areas in the north to non-pastoral areas in the south. Such a rapidly increasing and spreading epidemic trend deserved much attention. Though the government has noticed the problem and some countermeasures have already been taken (e.g. the application of brucellosis vaccine and the setup of brucellosis prevention institutes) [8–9], the situation was not optimistic yet.

Many studies have tried to explore reasons behind the prevalence, and there were some widely-accepted explanations [10], including the rapid development of husbandry, changes in the feeding mode of livestock, the frequent trading of livestock products among different areas, the increasing mobility of infected animals and so forth. In terms of the expansion of involved areas, some studies [11] assumed that people’s increasing opportunities to contact infected animals directly or indirectly in recent years might be the reason. In the study of Jiang et al [12], it was found that the Northern and Southern brucella strains shared the same MLVA-16 genotype, which somehow verified another popular speculation that the epidemic in southern area was partly caused by the import of infected animals from other areas. Therefore, it could be summarized that the development of husbandry and trading was the most commonly accepted explanation for human brucellosis epidemics in recent years.

Although current studies were instructive and illuminating for the prevention and control of human brucellosis, they still had some insufficiencies. First of all, most studies chose the qualitative methods instead of quantitative ones, which unavoidably made their conclusions rather crude and of less help. Moreover, some quantitative studies [13–14] have only focused on a limited geographic area or a particularly small population, which weakened their ability in providing evidence for carrying out more effective brucellosis countermeasures on a larger scale.

Another imperative need for human brucellosis study was to comprehensively consider the multivariate influences underlying the epidemics. The changing and not well-controlled epidemic of human brucellosis in recent years reminded us that there might be many factors interacting with each other and jointly influencing the incidence. However, some studies [6] only described the temporal and spatial characteristics of human brucellosis, but did not involve model building for further quantitative analysis (such as associated factors exploration and forecasting); some [15] built the time series model of human brucellosis incidence without penetrating its associated factors; another studies [16] considered the associated factors but limited to only a few aspects (such as environmental and animal husbandry factors). Hence, studies with comprehensive analysis of various associated factors are needed, especially considering the complex interaction among various factors. For example, suppose Cause A was the risk factor of human brucellosis, but it may be ignored in practice if there was another factor (Cause B) that offset the effect of Cause A on human brucellosis incidence. Therefore, it was necessary to consider the comprehensive effects of different causes all together in order to avoid bias and make the results of study more helpful and feasible.

This is a preliminary study targeted at the temporal-spatial characteristics of human brucellosis prevalence and the joint influence of different associated factors in mainland China in the last decade (2005 to 2016). We aimed to reveal the spatial and temporal characteristics of human brucellosis prevalence and explore the effects of its associated factors in a comprehensive and quantitative way, which would provide more valid evidence for making sophisticated and specific strategies for the prevention and control of human brucellosis in China and the rest of the world.

## Materials and Methods

### Materials

Incidence rates of human brucellosis of the 31 provinces in mainland China from 2005 to 2016 were excerpted from China Public Health Statistical Yearbook (http://www.nhfpc.gov.cn/zwgkzt/tinj/list.shtml) and the China National Knowledge Infrastructure (CNKI) website. A total of 13 associated factors were included and divided into three types: (1)Type I: the economy and animal husbandry factors (total output value of animal husbandry(*animal_husbandry*), amount of sheep (*sheep_num*), number of cattle (*cattle_num*), mutton production (*mutton_prod*), beef production (*beef_prod*) and Gross Domestic Product (*GDP*).);(2)Type II: the transportation factors (the turnover value of the whole society (*good_transfer*) and total length of highways (*highway*));(3)Type III: the hygiene and health factors (the number of medical institutions (*institute_num*), the number of health personnel (*health_personnel*), public health expenditure (*health_input*), urban medical expenditure proportion (*urban_medical_prop*) and rural medical expenditure proportion (*rural_medical_prop*). The corresponding data of associated factors came from the China Statistical Yearbook (http://www.stats.gov.cn/tjsi/ndsi/). For better understanding, abbreviations and meanings of each included factor were listed in Table 1. Furthermore, for the convenience of analysis, comparison and interpretation, all these factors were standardized beforehand.

**Table 1.**
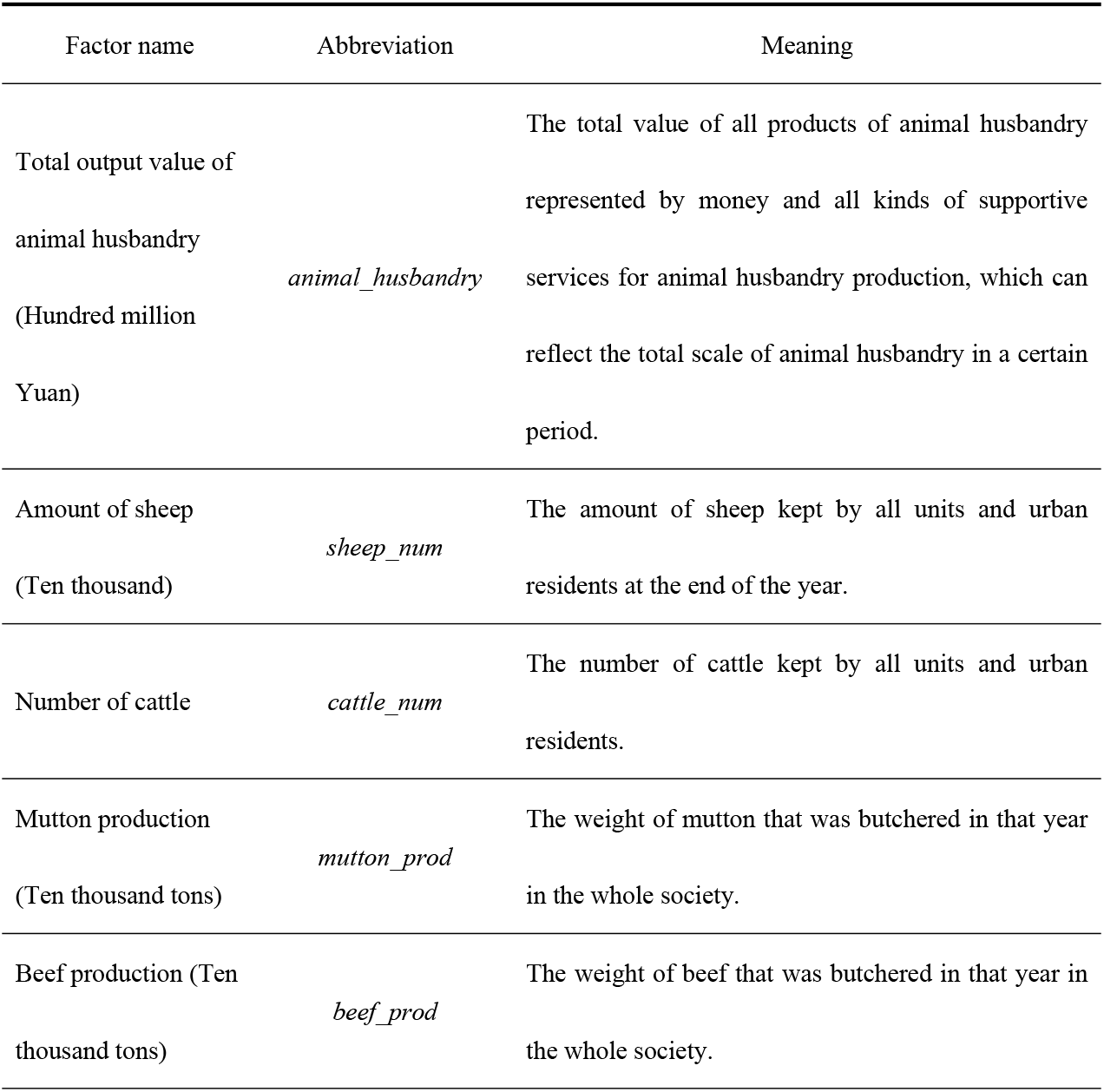
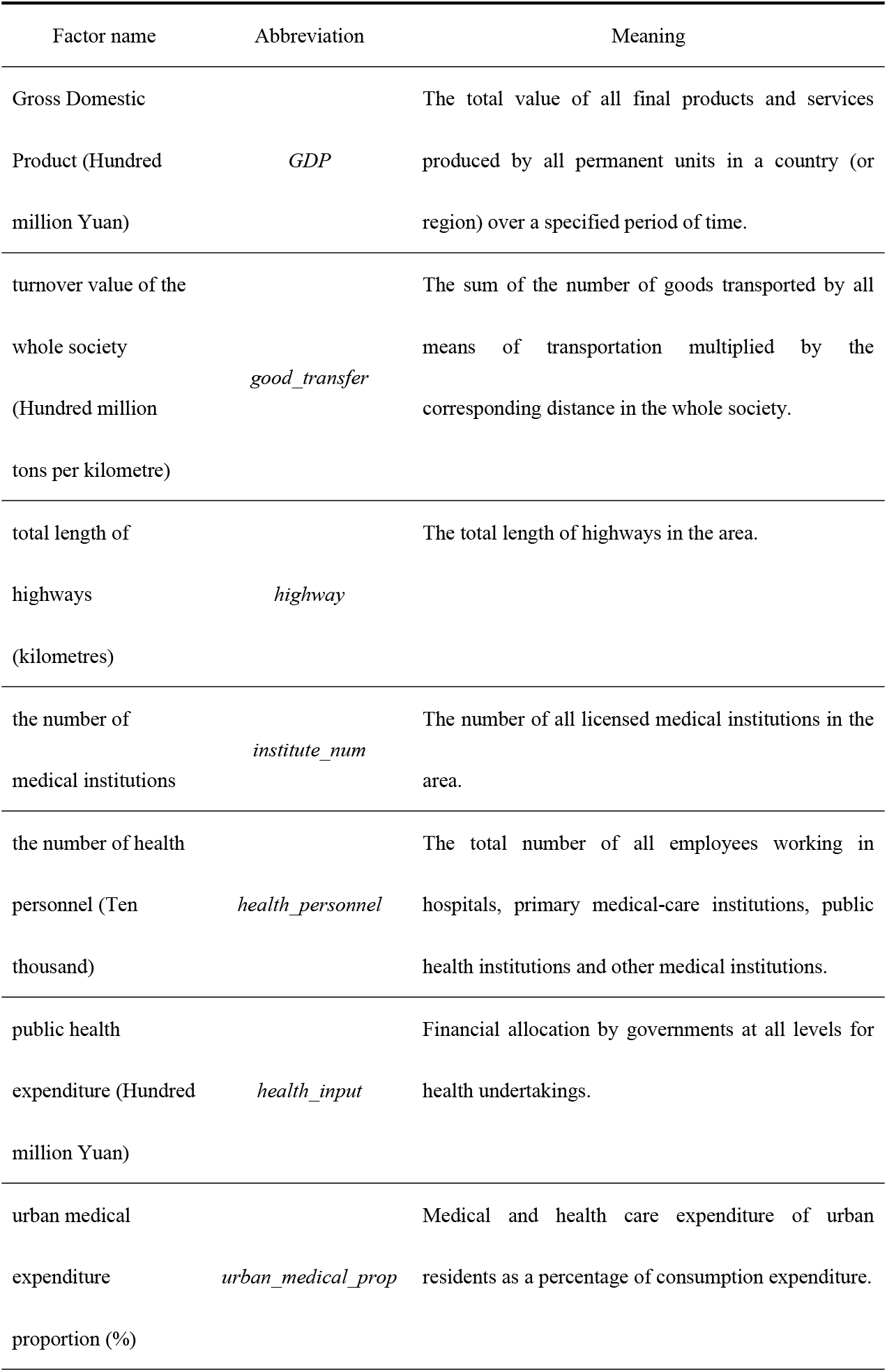
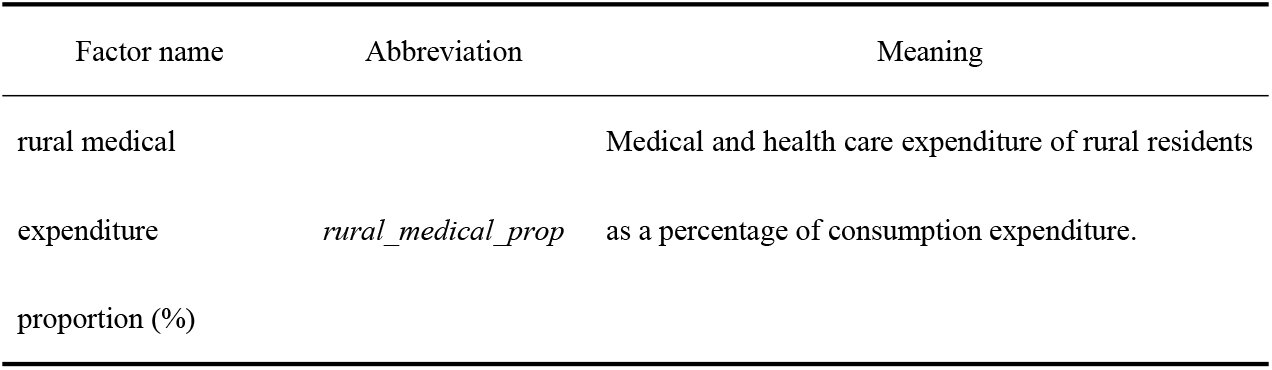
Explanations for all included associated factors.

| Factor name | Abbreviation | Meaning |
| --- | --- | --- |
| Total output value of animal husbandry (Hundred million Yuan) | *animal_husbandry* | The total value of all products of animal husbandry represented by money and all kinds of supportive services for animal husbandry production, which can reflect the total scale of animal husbandry in a certain period. |
| Amount of sheep (Ten thousand) | *sheep_num* | The amount of sheep kept by all units and urban residents at the end of the year. |
| Number of cattle | *cattle_num* | The number of cattle kept by all units and urban residents. |
| Mutton production (Ten thousand tons) | *mutton_prod* | The weight of mutton that was butchered in that year in the whole society. |
| Beef production (Ten thousand tons) | *beef_prod* | The weight of beef that was butchered in that year in the whole society. |
| Gross Domestic Product (Hundred million Yuan) | *GDP* | The total value of all final products and services produced by all permanent units in a country (or region) over a specified period of time. |
| turnover value of the whole society (Hundred million tons per kilometre) | *good_transfer* | The sum of the number of goods transported by all means of transportation multiplied by the corresponding distance in the whole society. |
| total length of highways (kilometres) | *highway* | The total length of highways in the area. |
| the number of medical institutions | *institute_num* | The number of all licensed medical institutions in the area. |
| the number of health personnel (Ten thousand) | *health_personnel* | The total number of all employees working in hospitals, primary medical-care institutions, public health institutions and other medical institutions. |
| public health expenditure (Hundred million Yuan) | *health_input* | Financial allocation by governments at all levels for health undertakings. |
| urban medical expenditure proportion (%) | *urban_medical_prop* | Medical and health care expenditure of urban residents as a percentage of consumption expenditure. |
| rural medical expenditure proportion (%) | *rural_medical_prop* | Medical and health care expenditure of rural residents as a percentage of consumption expenditure. |

### Methods

The temporal and spatial characteristics of human brucellosis epidemics from 2005 to 2016 in mainland China were described at first. Afterwards, cluster analysis was used to categorize the geographic regions based on time series incidence data of each province. As for the following exploratory analysis, it should first be noted that the complicated temporal-spatial effect within our dataset had violated the necessary independence assumption for the application of traditional linear regression models, while the sample size (observations of each province) was relatively small. Hence, to tackle these two challenges, this study intended to utilize the dynamic panel data model combined with supervised principal component analysis (PCA) to quantify the effects of associated factors on human brucellosis incidence. The basic form of dynamic panel data model (without supervised PCA) was shown in Eq.(1):

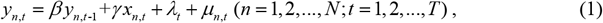

where *n* represented the province, *t* the year (*n* and *t* were both sequentially numbered), *y_n,t_* the human brucellosis incidence rate of the *n*-th province in the *t*-th year, and *y*_*n,t*-1_ referred to that in the past year. In addition, *x_n,t_* was the value of the associated factor of the *n*-th province in year *t, λ_t_* indicated the temporal effect of the year *t* and *μ_n,t_* the random effect of the *n*-th province in the *t*-th year. Specifically, the regression coefficient *γ* measured the relative risk of factor *x* on the human brucellosis incidence, which was the parameter of interest in this study.

During this study, we combined the supervised PCA with the dynamic panel data model [17]. Factors of concern were first selected based on their standardized univariate regression coefficients and the corresponding threshold *θ* estimated by cross-validation method. Subsequently, for each type of factors, we calculated the principal components as the linear combinations of those selected factors, where the linear combination coefficients were computed by the eigenvector-based method. For example, *economic*_1_, *economic*_2_,…, and *economic_R_* were used to represent the principal components of the type I factors (the economy and animal husbandry factors), where the value of *R* was determined by the cumulative contribution rate of the corresponding principal components. The cumulative contribution rate ranged from 0 to 100%, and the higher it was, the more eligible the principal components would be to represent those original factors. For type I factors, the cut-off point of the cumulative contribution rate was set to be 90%, which meant the *R* principal components of the type I factors should at least contain 90% of original information. Similarly, let *S* and *M* be the number of principal components for the type II factors (the transportation factors) and type III factors (the hygiene and health factors), respectively. The corresponding principal components can be denoted as *transfer*_1_, *transfer*_2_,…, and *transfer_S_*, as well as *health*_1_, *health*_2_,…, and *health_M_*. The cut-off point of the cumulative contribution rate was set to be 90% for the type II factors, and 80% for the type III factors (since only two original factors were included). As a result, the specific dynamic panel data model with the supervised PCA for this study could be built as below:

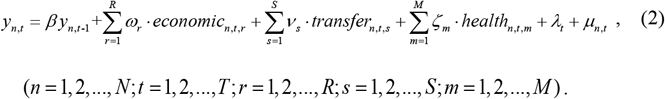

In Eq.(2), *economic_n,t,r_* referred to the value of the *r* principal component in economic and husbandry of the *n*-th province in the *t*-th year, so were the definition of *transfer_n,t,s_* and *health_n,t,m_*. In addition, *ω_r_, v_s_* and *ζ_m_* referred to the average value of the effect of the *r*-th principal component in economic and husbandry, the *s*-th principal component in transportation and the *m*-th principal component in hygiene and health, respectively. Definitions of other factors were the same as those in Eq. (1).

Throughout this study, all analyses were done in R 3.5.0, a free software environment for statistical computing and graphics. Computing Packages {*plm*} and {*ggplot2*} were downloaded from the Comprehensive R Archive Network (CRAN) at http://cran.r-project.org/ and installed in advance.

## Results

### Descriptions of the spatial and temporal distribution

According to the time series plot of the nationwide human brucellosis incidence (Fig 1), the incidence rate increased three-fold from 1.41 per 100,000 people in 2005 to 4.22 per 100,000 in 2016, though it went down a little in 2015. With such an upward trend nationally, the epidemic situation also changed slightly in different regions. Though the northern incidence rate has always been higher than that in the south, which was in accordance with previous reports [18], the southern incidence also began to increase in the recent five years and such an increase even continued despite the decrease in the northern incidence since 2014. From the time series plot, it could be inferred that: ① The human brucellosis incidence rate went up significantly during the study period; ②There was an overall upward trend of the epidemic in the south area (especially in southeastern provinces) while the northern provinces still kept high records of incidence rates. This indicated that the pastoral areas were still high epidemic areas while the incidence of human brucellosis also became more intense in half pastoral areas and agricultural areas, which coincided with the conclusion of Zhang et al [19].

**Fig 1.**
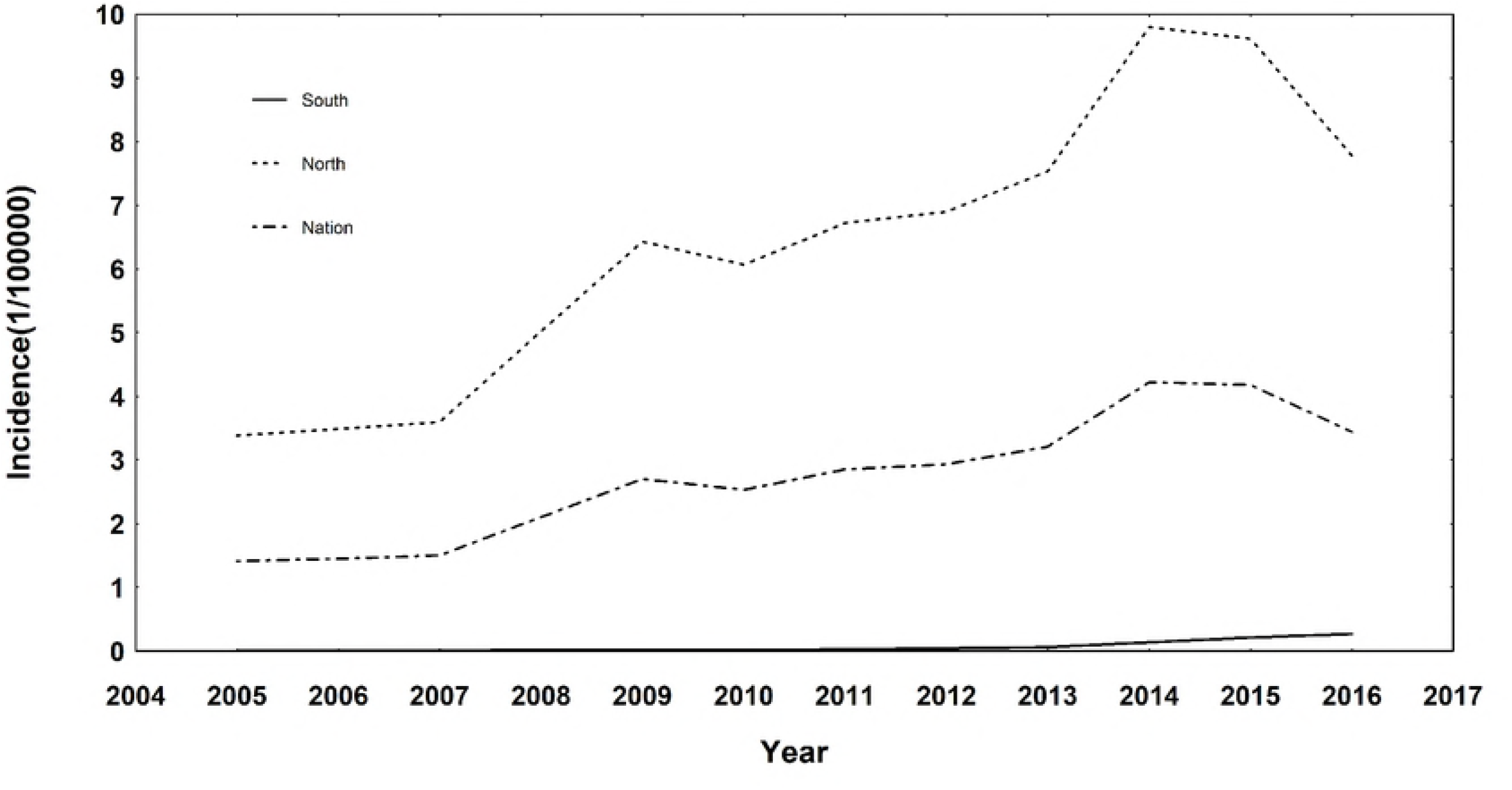
The time series plot of human brucellosis incidence nationwide and in south/north China.

As for the cluster analysis, the result indicated that these 31 provinces could be classified into three clusters according to their incidence level and epidemic trend. Specifically, Inner Mongolia alone belonged to Cluster 1; Ningxia, Xinjiang, Shanxi and Heilongjiang belonged to Cluster 2 and the remaining provinces were included in Cluster 3 (Fig 2). Through reviews of previous studies [6], a similar partition was observed, which helped to testify the rationality of our clustering.

**Fig 2.**
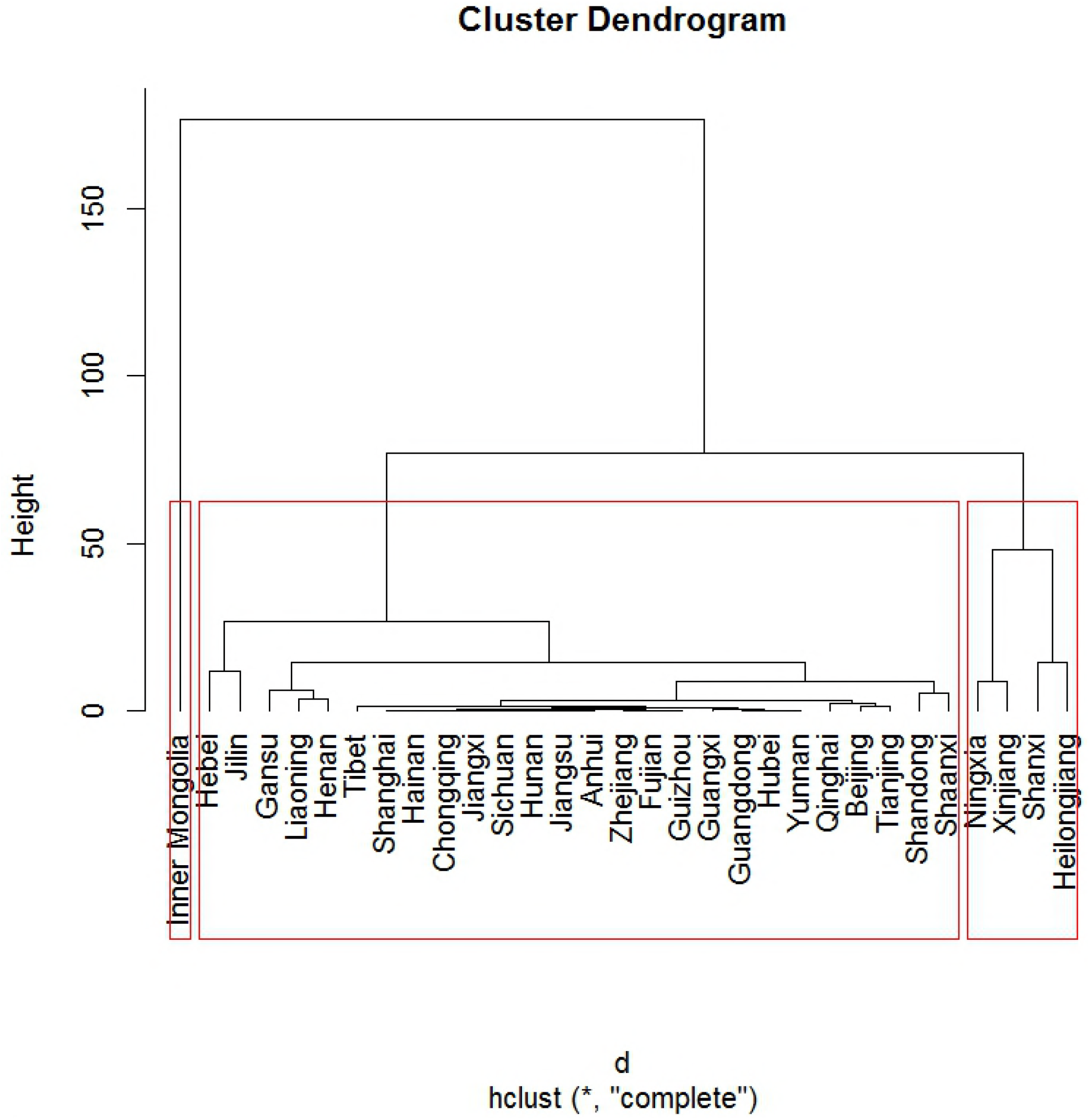
The clustering result of the 31 provinces.

Based on the clustering results, the time series plots for the three clusters were drawn respectively in Fig 3, which indicated that provinces in the same cluster did share similar prevalence level and epidemic trend. There was only one province in Cluster 1 (Inner Mongolia), and its incidence rate has always been the highest in China with a steady increase since 2005; after peaking at 2011, the rate dropped gradually despite a rebound in 2014. Ningxia, Xinjiang, Shanxi and Heilongjiang were categorized into Cluster 2 and their incidence rates were lower than that of Cluster 1, yet higher than most provinces in Cluster 3. Among these four provinces, Ningxia and Xinjiang shared a more similar epidemic characteristics (peaking at 2015 after increasing steeply since 2011) while incidence rates of Shanxi and Heilongjiang steadily remained at the level of 5-20 per 100,000 people most of the time. Cluster 3 included 26 provinces such as Jilin, Tibet and Guangdong. These provinces had low incidence rates and mostly peaked in 2014-2015, but Jilin was an exception for it experienced a decrease after an obvious rise in 2009.

**Fig 3.**
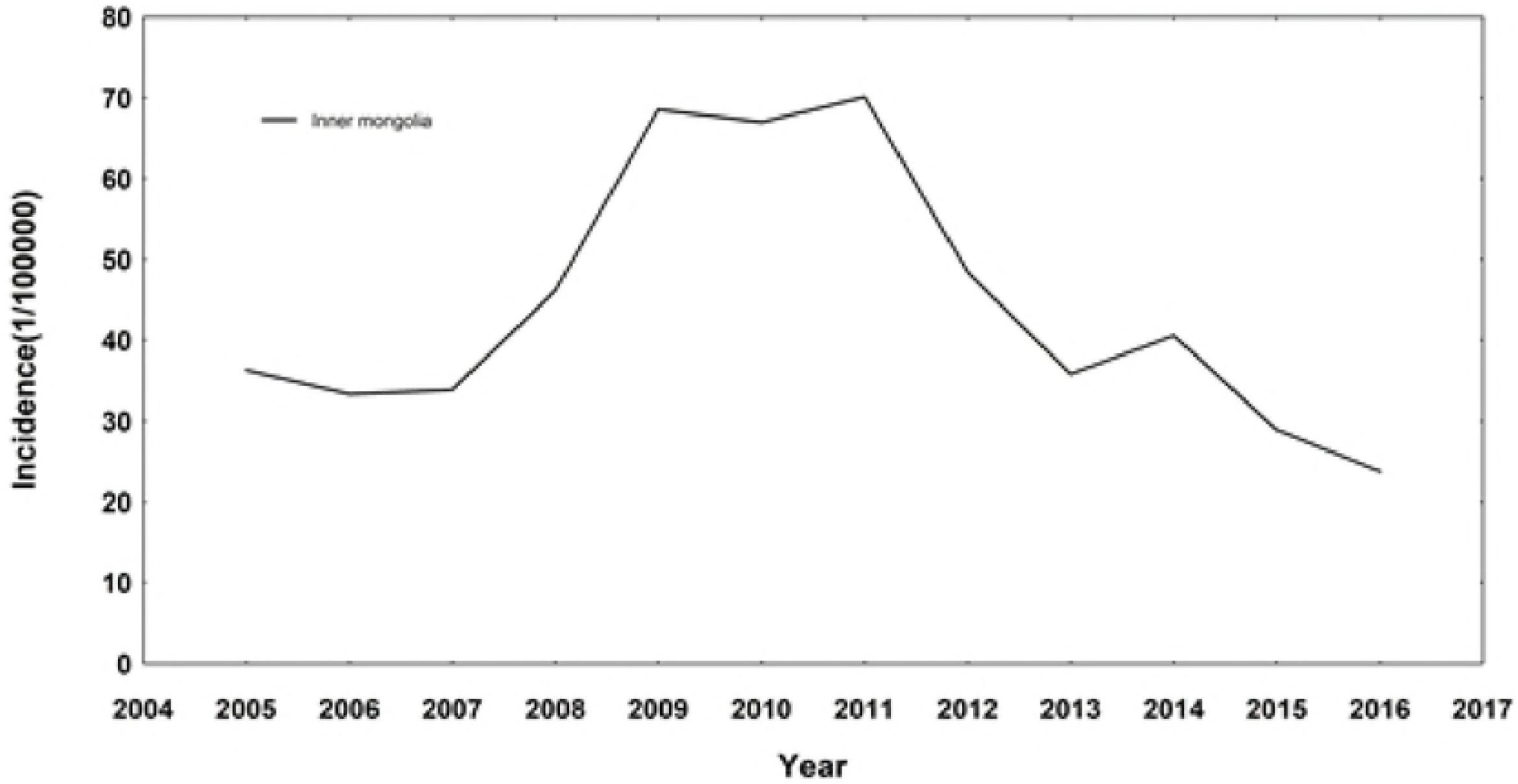
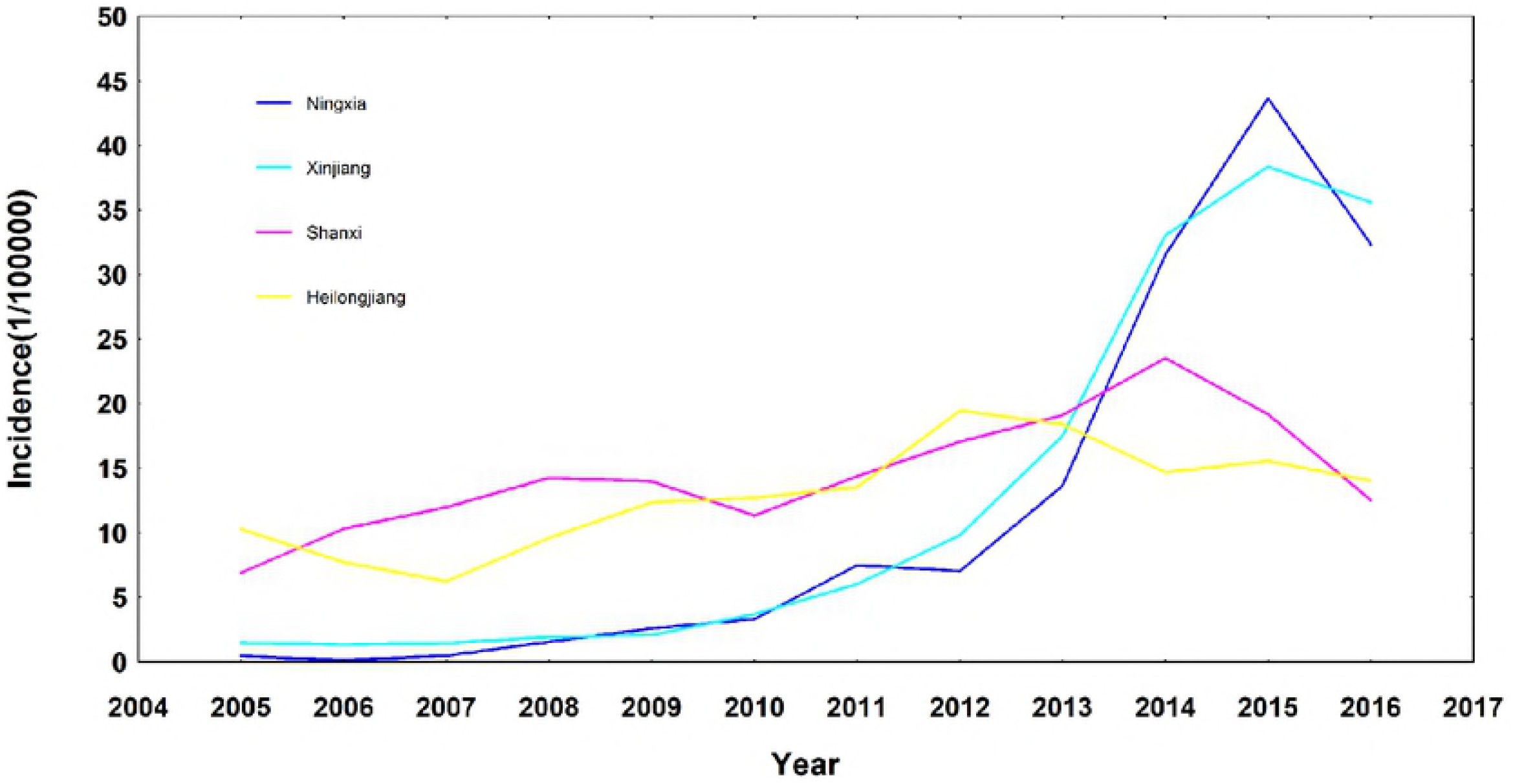
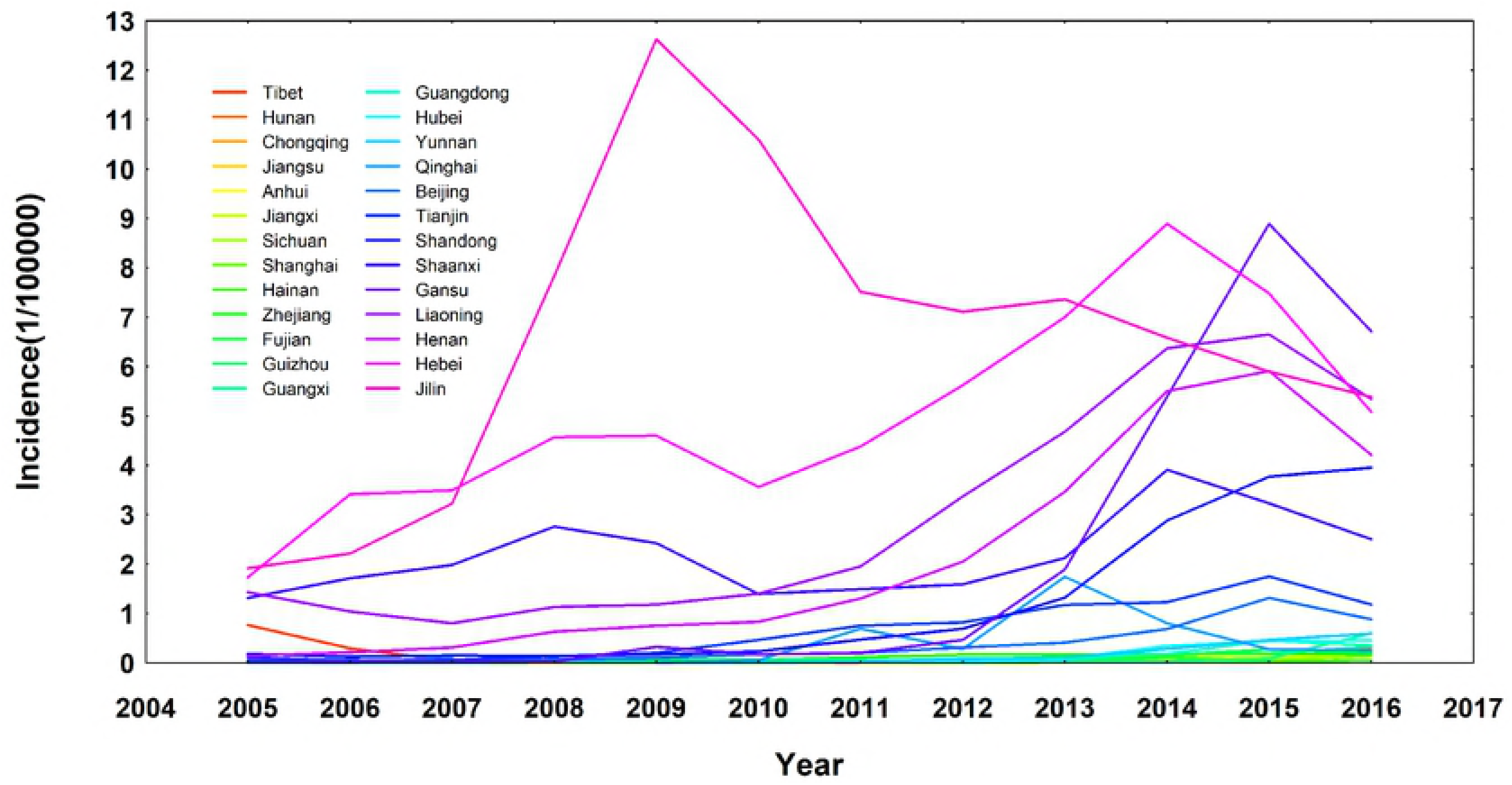
The time series plots of incidence rates for: (A)Cluster 1; (B)Cluster 2; (C)Cluster 3.

It could be seen that provinces in Cluster 2 (Shanxi, Heilongjiang, Ningxia and Xinjiang) and some provinces in Cluster 3 (Hebei, Liaoning, Shandong, Henan, Shaanxi and Gansu) all possessed a relatively high incidence rate and a similar epidemic trend, making it plausible to include them into further modelling analysis to reveal the underlying associated factors of human brucellosis. Other provinces were excluded because their extremely high or low incidence rates would perform as outliers and affect the validity and stability of statistical modelling. As a result, ten provinces in total were included in the statistical modelling stage, which were Gansu, Hebei, Heilongjiang, Liaoning, Henan, Ningxia, Shandong, Shanxi, Shaanxi and Xinjiang.

## Statistical modelling

Results of the dynamic panel model with supervised PCA involved two parts: one was the determination of principal components, and the other was the estimation and interpretation of associated factors’ effects on human brucellosis.

### Determination of principal components

Through calculation and comparison, 10 out of the 13 associated factors were selected as important associated factors, which were *GDP, animal_husbandry, sheep_num, mutton_prod, good_transfer, highway, institute_num, health_personnel, health_input* and *rural_medical_prop*. The following principal components were then formed using these selected factors.

(1) Principal components of the type I factors

The first two principal components were selected as representatives of the type I factors since their cumulative contribution rate was as high as 96.30%, and they were notated as *economic*_1_ and *economic*_2_ with the specific forms in Eq.(3) and (4).

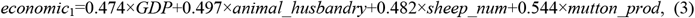

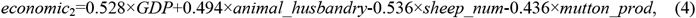

Since all coefficients in Eq.(3) were positive, *economic*_1_ could be considered as the general representative of the type I factors. In *economic*_2_, the coefficients of two factors (*GDP* and *animal_husbandry*) were positive while those of the other two factors (*sheep_num* and *mutton_prod*) were negative. Therefore, *economic*_2_ implied that the effects of *GDP* and *animal_husbandry* on human brucellosis might be different from those of the *sheep_num* and *mutton_prod*; in other words, *GDP* and *animal_husbandry* were more likely to be risk factors of human brucellosis.

(2) Principal components of the type II factors

Only the first principal component was selected to represent the type II factors, of which the cumulative contribution rate (84.96%) exceeded the threshold. This principal component was notated as *transfer* with the form of Eq.(5):

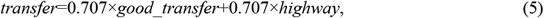

Coefficients in Eq.(5) were all positive, which indicated that *good_transfer* and *highway* might influence the human brucellosis incidence in a similar way.

(3) Principal components of the type III factors

The first two principal components were selected as representatives of type III factors since their cumulative contribution rate was 94.11%. They were notated as *health*_1_ and *health*_2_ with the form shown in Eq.(6) and (7).

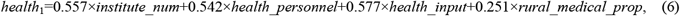

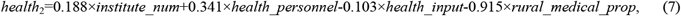

Again, all coefficients in Eq.(6) were positive, which meant that *health*_1_ could be taken as the general representative of type III factors. In *health*_2_, two coefficients (*institute_num* and *health_personnel*) were positive and two (*health_input* and *rural_medical_prop*) were negative, indicating that effects of *institute_num* and *health_personnel* on human brucellosis might not be the same as those of *health_input* and *rural_medical_prop*, and that *health_input* and *rural_medical_prop* were prone to be protective factors of human brucellosis.

### Estimation and interpretation of the associated factors’ effects

(1) The estimation results

The dynamic panel data model was built by including aforementioned principal components and the incidence rate in the past year (reflecting the dynamic characteristics of the model) as associated factors in it. Table 2 and Fig 4 showed the estimated coefficients and the goodness-of-fit results, respectively.

**Table 2.** The estimated coefficients of dynamic panel data model.

| Associated factor | Coef | SE | *t* | *P* |
| --- | --- | --- | --- | --- |
| $brucellosis_{t-1}$ | 0.95 | 0.04 | 22.31 | <0.001 |
| $economic_1$ | 0.65 | 0.35 | 1.88 | 0.06 |
| $economic_2$ | 0.40 | 0.42 | 0.96 | 0.34 |
| $health_1$ | - 1.14 | 0.47 | -2.42 | 0.02 |
| $health_2$ | 0.82 | 0.42 | 1.96 | 0.05 |
| $transfer$ | -0.87 | 0.47 | -1.86 | 0.07 |

**Fig 4.**
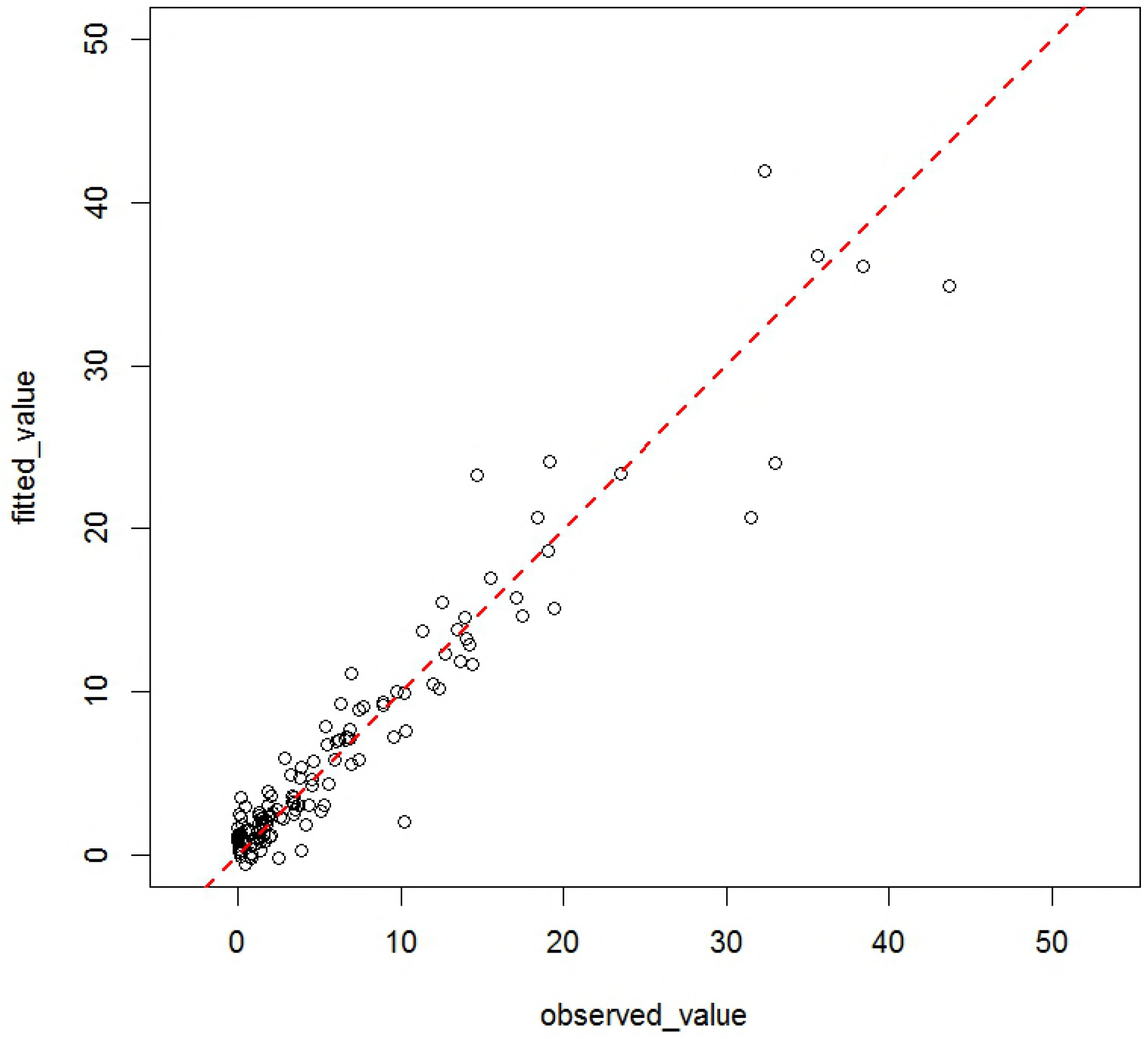
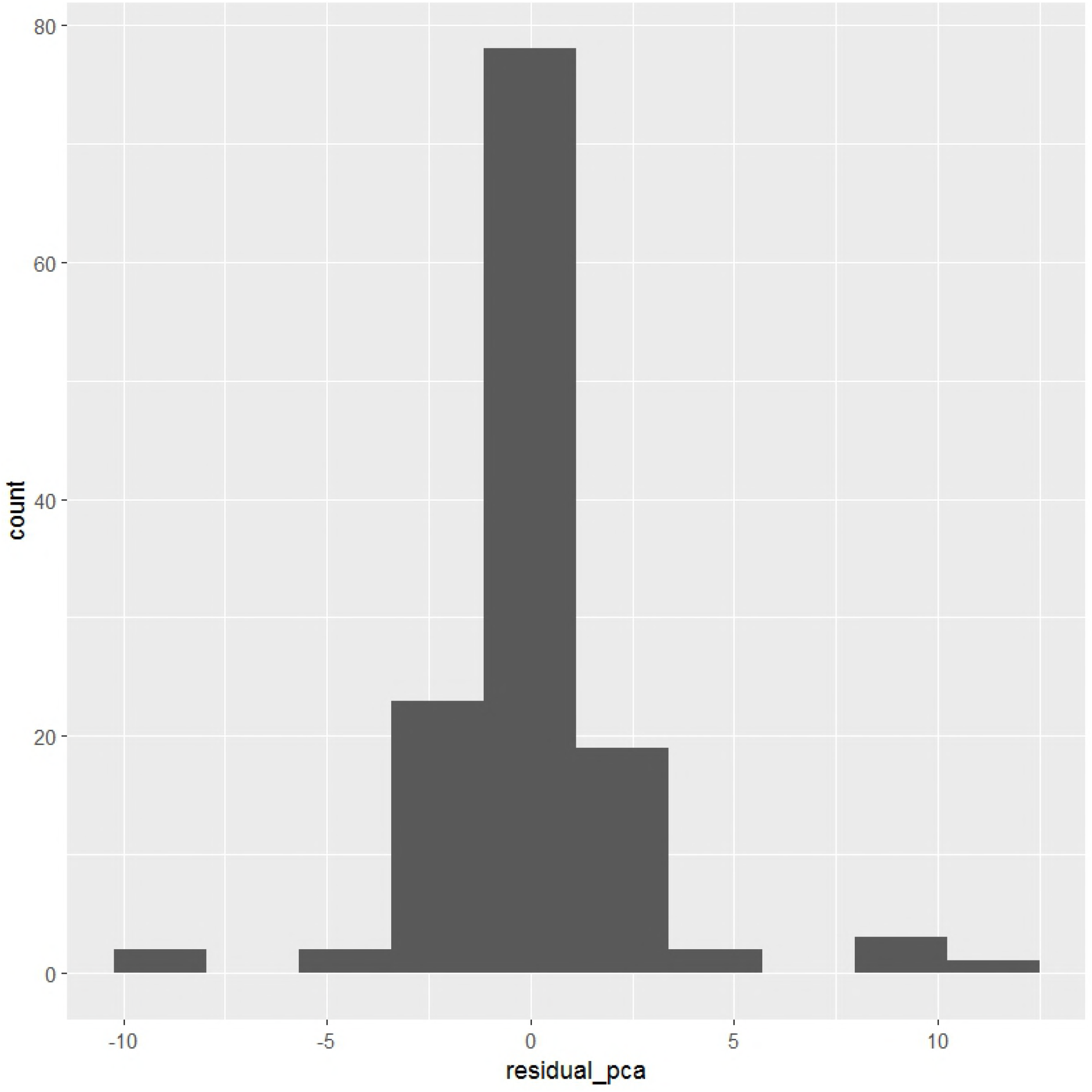
(A)The scatter plots of the observed incidence rates and the fitted values by model; (B)The histograms of the residuals of model.

According to Table 2, *brucellosis*_*t*-1_, *health*_1_ and *health*_2_ were associated with the current incidence of human brucellosis with statistical significance (P<0.05). Among these three factors, the coefficients of *brucellosis*_*t*-1_ and *health*_2_ were positive, while the coefficient of *health*_1_ was negative. Such results indicated that: ①The human brucellosis incidence in the previous year was positively related to the epidemic in the current year in the same place, which reflected the historical baseline effect of infectious diseases. ②A negative association existed between the overall health factors and human brucellosis, which meant the improvement of hygienic and health situations would possibly reduce the transmission risk of infectious diseases including but not limited to human brucellosis. In terms of *health*_2_, although the coefficient itself was positive, it should be reminded the coefficients of *health_input* and *rural_medical_prop* were negative according to Eq.(7). Hence, *health*_2_ emphasized that health input and rural medical expenditure proportion might play important roles in reducing the risk of human brucellosis.

In terms of the goodness-of-fit results, the corrected *R*^2^ of the model was 86.09%, which meant the model could explain more than 85% information of these factors’ effects on human brucellosis incidence. In addition, Fig 4A presented the comparison of the actual incidence rates and the fitted ones by the model. It could be seen that both the actual and fitted values situated on the diagonal line, which demonstrated that the two values were approximately the same. Besides, Fig 4B verified that the residuals of the model were normally distributed around zero, indicating that the model has sufficiently extracted information from data. Therefore, it was reasonable to conclude that the dynamic panel model with supervised PCA could appropriately clarify the effects of associated factors on human brucellosis.

(2) The interpretation of results

The estimated results in Table 2 could be further explained by inserting Eq.(3)~(7) into Eq.(2), which could be rewritten as Eq.(8) in a clearer way.

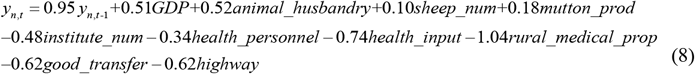

Because all factors in Eq.(8) have been standardized beforehand, the coefficient of each factor in this equation can be interpreted as the average changing value of human brucellosis incidence rate as the associated factor changes per standard deviation unit while the other factors remain constant. From the perspective of epidemiology, the average changing value can also be seen as the attributive risk (AR). Therefore, it could be concluded that GDP, total output value of animal husbandry, amount of sheep and mutton production were potential risk factors of human brucellosis while the others were potential protective factors. It was also worth noting that the absolute values of the coefficients of rural medical expenditure proportion (*rural_medical_prop*) and public health expenditure (*health_input*) were the largest two except *y*_*n,t*-1_, suggesting that these two might be the key points in the control of human brucellosis in China. More details of the effects on these associated factors would be discussed in the next section.

## Discussion

Human brucellosis is one of the few infectious diseases whose incidence rates still keep increasing nowadays in mainland China [20]. Though previous studies have tried to explore its epidemic patterns and associated factors, this study contributed to the prevention of human brucellosis in a more novel and more explicit way: it not only considered both temporal and spatial patterns of human brucellosis across mainland China, but also comprehensively revealed the multiple relations between human brucellosis and its potential associated factors. Therefore, the results of this study could supply the following new knowledge and implications for the prevention and control work in this field.

Firstly, apart from the traditional recognition that human brucellosis incidence rates in northern China were much higher than those in southern area, this study further analysed and compared the temporal and spatial epidemic characteristics of various areas. Specifically, according to both the incidence level and epidemic trend, all provinces of mainland China were classified into three clusters, i.e., Cluster 1 with the highest incidence rate all year round, Cluster 2 with rather high incidence rates but lower than that of Cluster 1 and lastly, Cluster 3 whose incidence rates were at a comparatively low level.

Secondly, this study jointly considered some potential associated factors of human brucellosis which used to be considered separately in previous studies. Results indicated that the human brucellosis incidence rate in the prior year may be a risk factor for the current year’s possible epidemic (AR=0.95). More importantly, results also implied that public health expenditure and rural medical expenditure proportion might be two important protective factors of human brucellosis (AR =−0.74 and −1.04 respectively).

Based on these new results, this study discovered that the practical work of human brucellosis prevention and control could be improved in at least two ways:

(1) Clustering could be used to better reveal the heterogeneity and complexity of the transmission dynamics of human brucellosis in different spatial and temporal settings. This study classified all provinces in mainland China into three clusters depending on the similar incidence level and epidemic trend within a cluster. Such partition could provide evidence for more effective countermeasures with greater pertinence targeting at various areas. Though some scholars [21] have previously tried to categorize areas in mainland China based on their incidence rates, this study put it further by considering both the incidence rate and the epidemic trend. By doing this, researchers could better reveal the association, know more about the epidemic mechanism as well as offer more practical clues for adjusting measures in different conditions.
(2) The exploratory analysis in this paper contributed to clarify the potential effects of associated factors on human brucellosis in a more comprehensive way. It helped update and deepen the understanding of epidemic mechanism as well as locate the key areas of controlling human brucellosis more precisely. Here are some brief discussions on some factors involved.

① **Historical impact** According to the analysis, there existed a statistically significant influence of the human brucellosis incidence rate in the previous year (lag, brucellosis 1) on the current incidence rate, which might be explained by the latency and invasiveness of brucella [22].
② **Transportation impact** Many scholars have assumed that the smuggling of infected animal products from other provinces might account for the human brucellosis epidemics in those newly-emerging areas. However, the transportation impact turned out to be insignificantly negative in this study. One potential reason was that this study chose the turnover value of the whole society (*good_transfer*) and total length of highway routes (*highway*) to represent transportation, but in real life, the smuggling of animal products (especially those infected ones) tended to depend more on hidden routes instead of highways. The other possible reason was that better transportation condition always associated with more advanced economic and social development, and in such case, the quarantine and inspection measures and regulations would be more sophisticated and stricter, which would reduce the smuggling of infected animal products.
③ **Animal husbandry impact** Another widely-accepted opinion was that the development of husbandry and the change in feeding pattern might be the cause of human brucellosis [19], but no supporting result was found in this study. This reminded us that though there was a possibility that the development of husbandry created a more suitable environment for the epidemic of human brucellosis, some other factors might minimize or even counteract such a risk. However, this result did not implicate to deny the influence of husbandry development on human brucellosis; on the contrary, it emphasized that the risk created by the development of husbandry could be reduced if much attention could be paid to some protective factors of human brucellosis.
④ **Health impact** One interesting point of this study was that hygiene and health condition was negatively associated with human brucellosis epidemics. It suggested that efforts in improving the quality of community- or village-level public health services might bring unexpected benefits to the work of human brucellosis prevention and control. To be more specific, increasing the public health expenditure, especially the rural medical expenditure proportion might be a potential breakthrough for handling human brucellosis in the future.

The practical work of human brucellosis prevention and control involved allocating public resources in the most suitable and most cost-effective place especially when the budget is limited. To this end, combined with the results of this study, some following advices could be proposed: (I)Governments should further enhance the communication and corporation among hospitals, township health centres and rural clinics as well as increase the investment in economy and infrastructure in medical institutes [23–24]. Considering that the rural and traditional pastoral areas are still high-risk regions, it is recommended to pay more attention to the improvement of the quality of village-level public health service in the countryside, which helps to better carry out tertiary prevention among residents living there. (II)Residents’ initiatives in cooperating with the prevention work should be better encouraged, and this can be done by regular health education and propaganda. (III)The prevention work can also be improved by raising the health awareness of rural residents and perfecting the new rural cooperative medical system [25]. Governments can increase the subsidized expenditure of health care in rural areas so as to maximize the possibility of an increase of the rural medical expenditure proportion. Noticing that residents’ incomes can also affect their medical expenditure [26], current priority could be given to increase the health care subsidy of residents in these areas in order to achieve the goal of improving the medical expenditure proportion to control human brucellosis.

Outside mainland China, many other regions in the world also suffer from human brucellosis, such as Southern Brazil and Portugal [27–28]. Although China is a country large in land and diverse in population composition, which could enhance the value of this study in providing evidence for better control measures in other areas, it is possible that different countries may have different reasons for human brucellosis epidemics. On this basis, further studies with cross-national data and more associated factors are expected to contribute to faster and better prevention and control of human brucellosis worldwide.

## Acknowledgements

We gratefully thank Miss Minghan Xu for giving advice on the manuscript of this study.

